# Latitudinal Variation in Life History Reveals a Reproductive Disadvantage in the Texas Horned Lizard (*Phrynosoma cornutum*)

**DOI:** 10.1101/364091

**Authors:** Daniel F. Hughes, Walter E. Meshaka, Carl S. Lieb, Joseph H. K. Pechmann

## Abstract

Geographically widespread species that occupy many thermal environments provide testable models for understanding the evolution of life-history responses to latitude, yet studies that draw range-wide conclusions using descriptive data from populations in the core of a species’ distribution can overlook meaningful inter-population variation. The phrynosomatid lizard *Phrynosoma cornutum* spans an extensive latitudinal distribution in North America and has been well-studied in connection with life-history evolution, yet populations occupying the most northern and coldest areas within its range were absent from previous examinations. We tested genus-wide models and challenged species-specific findings on the evolution of the life-history strategy for *P. cornutum* using populations at the northern edge of its geographic range and comparative material from farther south. Multivariate analyses revealed that egg dimensions decreased with clutch size, suggestive of a previously unrecognized tradeoff between egg size and egg number in this species. Interestingly, reproductive traits of females with shelled eggs did not covary with latitude, yet we found that populations at the highest latitudes typified several traits of the genus and for the species, including a model for *Phrynosoma* of large clutches and delayed reproduction. A significant deviation from earlier findings is that we detected latitudinal variation in clutch size. This finding, although novel, is unsurprising given the smaller body sizes from northern populations and the positive relationship between clutch size and body size. Intriguing, however, was that the significant reduction in clutch size persisted when female body size was held constant, indicating a reproductive disadvantage to living at higher latitudes. We discuss the possible selective pressures that may have resulted in the diminishing returns on reproductive output at higher latitudes. Our findings highlight the type of insights in the study of life-history evolution that can be gained across Phrynosomatidae from the inclusion of populations representing latitudinal endpoints.

---

Nearly all animal behaviors are affected by latitudinal variation in climate, including metabolism (Tsuji, 1988), seasonal activity (Sperry et al., 2010), and predator-prey interactions (Laurila et al., 2008). These phenomena are intensified in ectotherms because temperature strongly influences their physiology (Huey, 1982), and many aspects of physiology have implications for demography (Huey and Berrigan, 2001). For example, growth is slower at higher latitudes from decreased temperatures that reduce activity and increase dormancy (e.g., Blouin-Demers et al., 2002; Laugen et al., 2003). Latitudinal variation in ectotherm body size, growth, and mortality has been well studied (e.g., Adolph and Porter 1993; 1996), however, we know far less about such variation from direct measures of reproduction. Changes in fecundity can have significant demographic costs that impact population size and persistence. For example, smaller ectothermic lizards generally have fewer young (Fitch, 1985). If lizards are smaller at higher latitudes (e.g., Ashton and Feldman, 2003), then clutch size is expected to be smaller. Nevertheless, there is a tradeoff between egg size and clutch size because the offspring number cannot be changed without changing egg size and thus offspring size and ultimately survival (Sinervo, 1990). Herein we examine variation in reproductive traits of Texas horned lizards (*Phrynosoma cornutum*) from three populations that span most of the species’ latitudinal distribution in the United States.

Body size in lizards is highly variable (Olalla-Tárraga et al., 2006), and patterns often do not conform to traditional latitudinal explanations for vertebrates (e.g., Bergmann’s Rule [Angilletta et al., 2004]). Clutch size is tightly linked to body size in lizards (King, 2000), so fecundity should vary with latitude. Although body size in lizards is constrained by seasonal resource acquisition (e.g., Ballinger, 1977; Van Noordwijk and de Jong, 1986), the amount of time each year that foraging is feasible will decline with increasing latitude. Consequently, latitudinal patterns in lizard body size covary with several life-history variables, one of which is clutch size (Angilletta et al., 2004). At higher latitudes, growth is slower as activity seasons become shorter; a relatively larger body size is usually achieved by prolonged growth and delayed maturation (Atkinson, 1994). It follows that clutch size should vary widely with latitude, largely reflecting variation in the active season. Further, latitudinal variation in clutch size should mirror the variation in body size, and any deviations from this pattern would suggest differential selective pressures within species populations (Tinkle et al., 1970).

North American horned lizards (Phrynosomatidae: *Phrynosoma*) have previously been the subject of many ecological studies in connection to life-history theory (e.g., Ballinger, 1974; Howard, 1974). Pianka and Parker (1975) conducted a genus-wide examination of horned lizard species, and described their life-history strategy as belonging to the “delayed reproduction, large clutch and single brooded” category per Tinkle et al. (1970). Further, Pianka and Parker (1975) sought to reconcile the heterogeneous life-history strategy in horned lizards (i.e., r-selected for high fecundity and K-selected for delayed maturity and long lives) with the expectation of higher juvenile mortality relative to the more homogenous strategies for other phrynosomatid lizards (e.g., Adolph and Porter, 1993; Pianka, 1970).

Studies of the Texas horned lizard (*P. cornutum*) have corroborated several genus-wide patterns and hinted at some geographic trends to its life-history strategy. Across a latitudinal gradient from Mexico to Colorado, Texas horned lizards exhibit variation in body size with smaller individuals at northern latitudes (Montgomery et al., 2003), yet not all horned lizards exhibit this trend (Pianka and Parker, 1975). The large clutch sizes and delayed maturity among horned lizards is associated with the production of a single clutch per season (Pianka and Parker, 1975). Clutch frequency in the Texas horned lizard, however, is geographically variable among populations without demonstrating a regional geographic bias (Ballinger, 1974; Howard, 1974; Vitt, 1977). A seasonal increase in testis size from May–August is conserved across horned lizards, and this increase is consistent across longitude in the Texas horned lizard (Ballinger, 1974; Howard, 1974). Delayed reproduction is a genus-wide phenomenon and occurs in Texas horned lizards from the southwestern United States (Howard, 1974) to Oklahoma (Endriss et al., 2007). Texas horned lizards exhibit latitudinal variation in clutch size from Texas to Oklahoma with smaller clutches observed in populations at higher latitudes (Howard, 1974; Vitt, 1977; Endriss et al., 2007). However, this latitudinal pattern is interpreted without accounting for established clinal changes in adult body size (i.e., Montgomery et al., 2003).

Museum specimens were the primary source of data for nearly all of the studies on the Texas horned lizard cited above. In keeping within this well-established framework, we examined a large series of museum specimens from the northern edge and a southern portion of the Texas horned lizard’s geographic range to address the following three main questions: (1) are there geographic patterns in life-history characteristics after controlling for differences in body size among populations? (2) what is the relationship between egg size and clutch size within and among populations? (3) are there latitudinal patterns in multivariate life history? Importantly, we included populations from a higher latitude than previously studied, and we focused on reproductive traits for which predictability has been lacking in previous investigations. Essentially, we attempted to use our data—that include the northernmost populations—to resolve previously inconsistent life-history findings deduced from narrow geographic sampling for the Texas horned lizard.

## MATERIALS AND METHODS

We examined 213 specimens of the Texas horned lizard (Fig. 1) from the herpetology collections of the Sternburg Museum (Hays, Kansas) and the University of Texas at El Paso Biodiversity Collections (El Paso, Texas). The specimens originated from three primary locations: Kansas, Texas, and New Mexico (Fig. 2). Eighty-seven specimens were collected during 1958–2013 from seven northern counties in Kansas (Ellis, Lincoln, Osborne, Rooks, Russell, Saline, and Trego). Seventy-three specimens were collected during 1958–2010 from three western counties in Texas (Culberson, El Paso, and Hudspeth). Forty-five specimens were collected during 1966–2014 from 10 mostly southern counties in New Mexico (Chaves, Doña Ana, Eddy, Grant, Hidalgo, Luna, Otero, San Miguel, Sierra, and Socorro). We also examined a smaller sample of eight specimens that were collected during 1962–1979 from three northern states in Mexico (Chihuahua, Coahuila, and Durango).

**Fig. 1.**
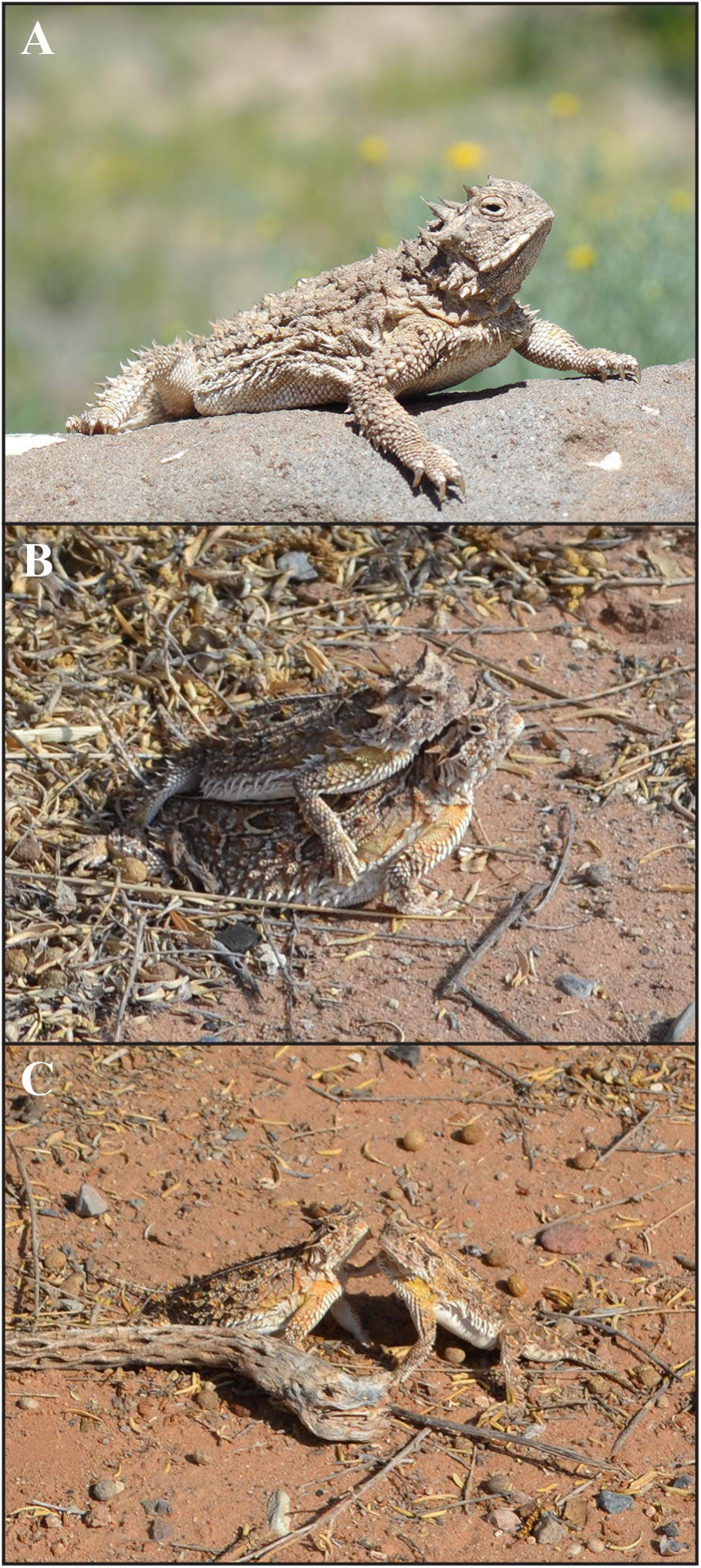
Texas horned lizards (*Phrynosoma cornutum*) from Dñna Ana County, New Mexico. (A) Gravid female, (B) male and female in coitus, and (C) facial rubbing immediately post coitus.

**Fig. 2.**
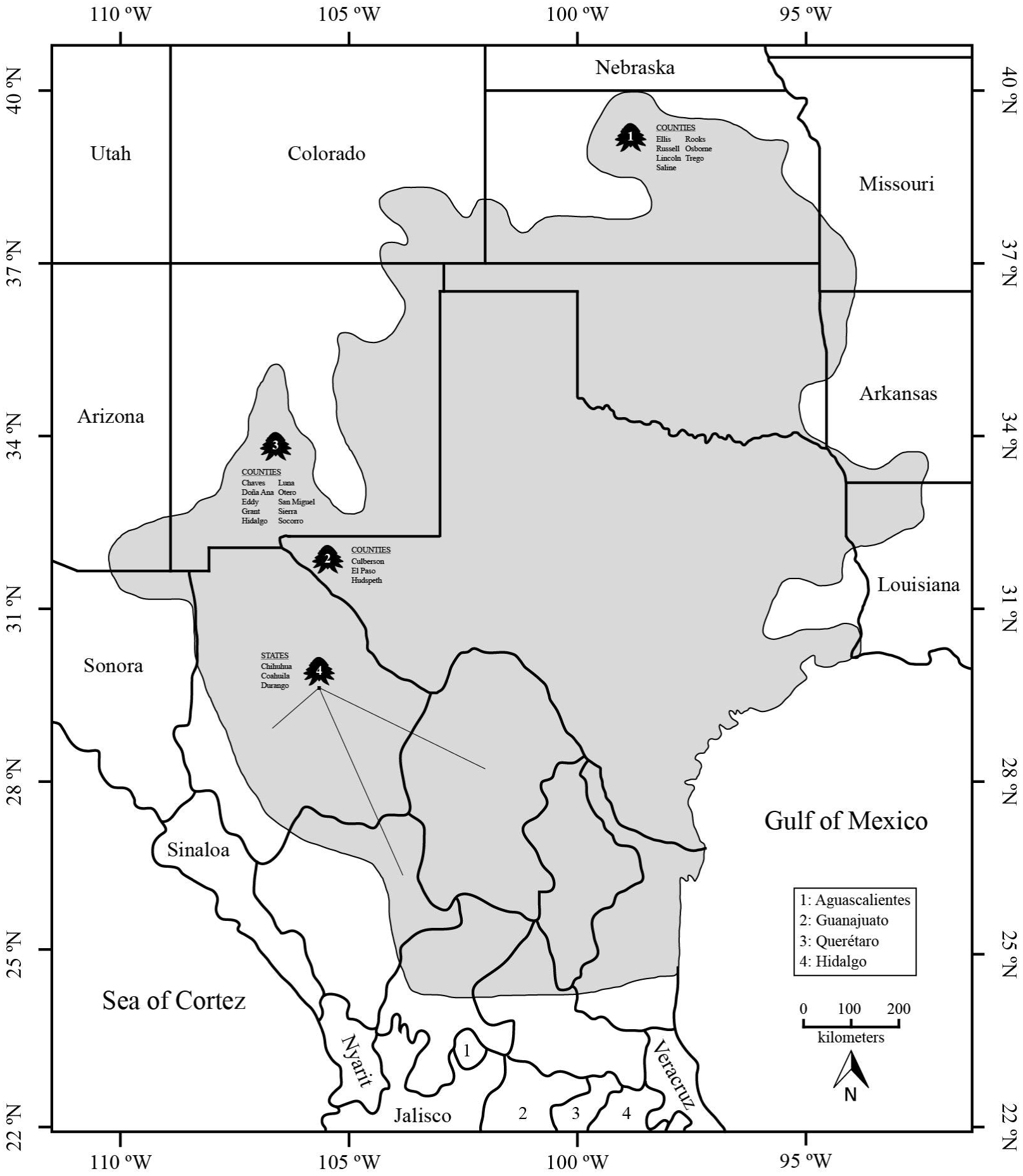
Map showing the populations of the Texas horned lizard (*Phrynosoma cornutum*) we examined for life histories: Kansas (1), Texas (2), New Mexico (3), and Mexico (4). The species’ geographic distribution (shaded area) was modified from Hammerson (2007).

For each specimen, we measured snout-vent length (SVL) with calipers to the nearest 0.1 mm. Lizards were then dissected and sexed. Females were classified as immature or non-reproductive if the largest ovarian follicle was < 3 mm and not vitellogenic. Females were assigned as reproductive if they contained shelled eggs or vitellogenic ovarian follicles ≥ 3 mm. Diameters of the largest yolking follicles, and length and width of shelled eggs were measured to the nearest 0.1 mm. Egg volumes were calculated from the equation for an ellipsoid: v = (4/3) πab^2^ (Mayhew, 1963). Clutch volumes were calculated as the product of clutch size and egg volume. The fraction of length and of mid-testis width/male SVL was used as a measure of seasonal male fertility. The ovarian cycle was determined by plotting monthly distribution of the largest follicle and shelled egg length.

Without direct observations on the number of eggs deposited in each clutch, we estimated clutch size using counts of shelled eggs and/or vitellogenic ovarian follicles ≥ 3 mm. We used a General Linear Model (GLM) with SVL as a covariate to test if there was a significant difference between the two clutch measures (shelled eggs and vitellogenic follicles). We found that little overlap existed in body sizes among populations (Fig. 3), and thus we tested whether the slopes of the regression lines between SVL and clutch size differed among populations or by clutch measure by testing for interaction effects among these variables using a GLM. Because SVL is linearly related to clutch size in Texas horned lizards (Ballinger, 1974; Howard, 1974) and to avoid potential problems with the use of ratios in statistics (Packard and Boardman, 1999), all measurements were analyzed in an analysis of covariance (ANCOVA) with SVL as a covariate to remove the effect of body size in our comparisons of life-history variables among populations.

**Fig. 3.**
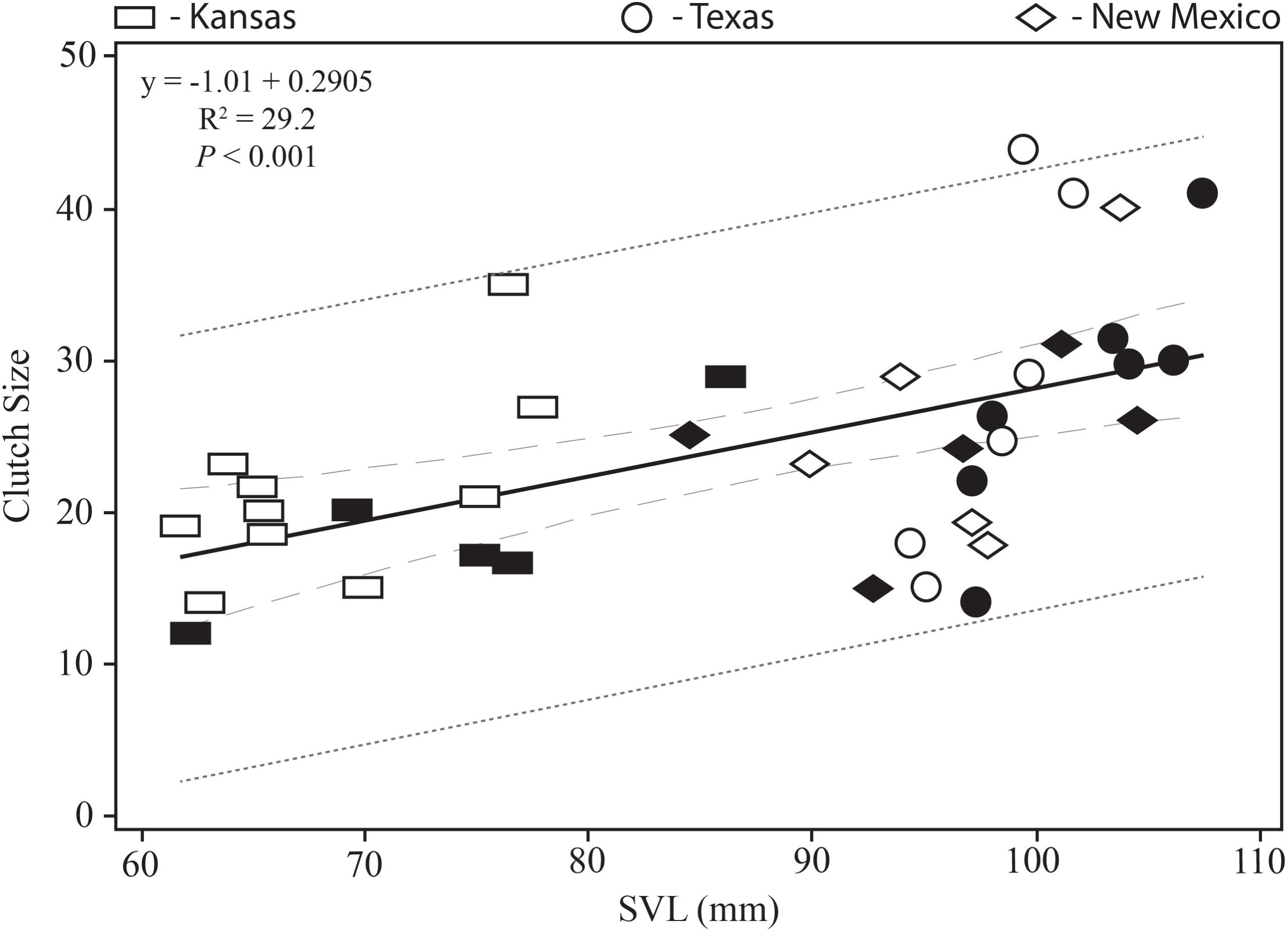
Clutch size regressed against body size for three populations of the Texas horned lizard (*Phrynosoma cornutum*). Filled symbols represent shelled eggs and open symbols vitellogenic ovarian follicles. The prediction interval is denoted by a dark-grey tightly spaced dashed line and the 95% confidence interval is denoted by a light-grey loosely spaced dashed line.

We explored multivariate divergence of the three primary populations with principal component analysis (PCA). To avoid weighting variables by their variance, the PCA was performed using a correlation matrix calculated from population means of five life-history characters (clutch size, egg length, egg width, egg volume, and clutch volume). We restricted this analysis to females with shelled eggs only. One female specimen from New Mexico with shelled eggs was excluded because of the eggs were rounded. Measurements were natural-log transformed and size corrected using SVL as a covariate because all five measurements were positively correlated with body size. The residuals were saved and included as input for PCA.

We used standardized guidelines to determine that our data met the assumptions (e.g., normality) of PCA (McGarigal et al., 2000). All statistical analyses were conducted in Minitab v. 17 (Minitab Statistical Software, State College, Pennsylvania, USA).

## RESULTS

### Testicular cycle

Testis length and width varied seasonally in all three populations (Fig. 4A–C). Among males in Kansas and Texas, testes were largest in April and smallest in July (Fig 4. A, B). In New Mexico, testes reached a maximum size in June and rapidly decreased in size thereafter, with the smallest sizes evident in July (Fig. 4C).

**Fig. 4.**
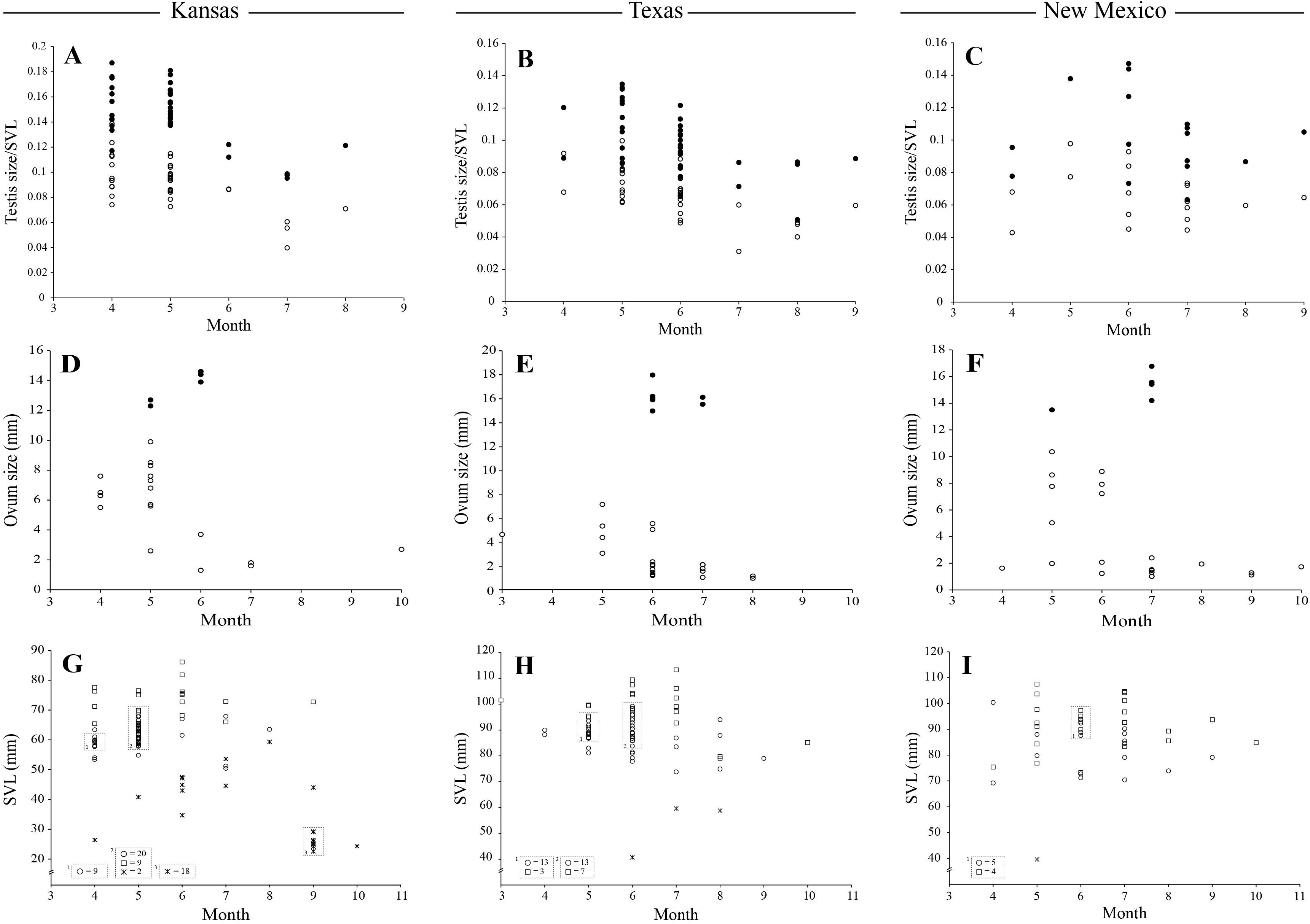
Monthly distribution of three life-history traits in the Texas horned lizard (*Phrynosoma cornutum*) from three populations. Top row: Testis length and width as a fraction of male snout-vent length (SVL) for (A) Kansas (n = 39), (B) Texas (n = 38), and (C) New Mexico (n = 17). Filled circles represent testis length/SVL and open circles testis width/SVL. Middle row: Largest ovarian follicle diameters and shelled egg lengths for (D) Kansas (n= 26), (E) Texas (n = 27), and (F) New Mexico (n = 25) New Mexico. Filled circles represent shelled eggs and open circles ovarian follicles. Bottom row: Body size (SVL) for (G) Kansas (39 males, 25 females, 22 juveniles), (H) Texas (42 males, 26 females, 3 juveniles), and (I) New Mexico (19 males, 25 females, 1 juvenile).

### Nesting season

The nesting season was short (ca. 2–3 months) in all three populations, and some overlap was evident among them (Fig. 4D–F). In Kansas, shelled eggs were present in females during May–June (Fig. 4D). In Texas, ovigerous females were present also for two months, beginning one month later (June–July) (Fig. 4E). The longest nesting season (May–July) was detected in New Mexico, which subsumed the combined nesting season of the other two regions (Fig. 4F).

### Clutch size

Because we had two measures of clutch size (shelled eggs and yolked ovarian follicles) from three discrete populations (Kansas, Texas, and New Mexico), we first tested for interactions between various variables associated to know if we could reliably compare clutch sizes among populations. The interaction between population X clutch measure was not significant (*F*_2,37_ = 0.51, *P* = 0.607), and the interaction among population X clutch measure X body size was also not significant (*F*_2,37_ = 0.46, *P* = 0.637). As a result, we felt that the intercept of these regression lines could be validly compared because the slopes are essentially parallel (Fig. 2). Further, we found that there was no difference in our clutch size estimates between the two measures of clutch size among the three populations (*F*_1,37_ = 3.41, *P* = 0.074) and thus we pooled these values for the following analyses involving clutch sizes. After removing the effects of maternal body size, we found that there was a significant effect of source population on clutch size, with the smallest mean clutch size found in Kansas, intermediate in New Mexico, and largest in Texas (*F*_2,37_ = 3.62, *P* = 0.039) (Table 1). Size-corrected analyses detected no significant difference among populations in mean clutch volume (*F*_2,15_ = 1.2, *P* = 0.336) (Table 1).

**Table 1.** Summary of life-history characters for populations of the Texas horned lizard (*Phrynosoma cornutum*). Means ± standard deviation are followed by ranges and samples sizes for body size, clutch size, egg length, egg width, egg volume, and clutch volume.

| Variable | Kansas | Texas | New Mexico | Mexico |
| --- | --- | --- | --- | --- |
| Male SVL (mm) | 60.5 ± 4.3 | 87.3 ± 5.3 | 83.6 ± 9.1 | 90 ± 6.4 |
|  | 50.4–68 | 73.8–99.2 | 69.2–100.4 | 83.5–98.1 |
|  | 39 | 43 | 19 | 5 |
| Female SVL (mm) | 70.8 ± 6.4 | 96.8 ± 8.5 | 92 ± 9.1 | 103.9 ± 5.9 |
|  | 61.6–86.1 | 79.1–113.3 | 73.2–107.5 | 99.8–108.1 |
|  | 26 | 27 | 25 | 2 |
| Juvenile SVL (mm) | 36.6 ± 11.9 | 53 ± 10.7 | 39.6 | 46.1 |
|  | 22.6–59.3 | 40.7–59.6 | 1 | 1 |
|  | 22 | 3 |  |  |
| Clutch Size (pooled) | 20.6 ± 6 | 28.2 ± 9.7 | 25 ± 7.2 | |
|  | 12–35 | 14–44 | 15–40 | - |
|  | 15 | 13 | 10 |  |
| Yolked Ovarian Follicles | 21.4 ± 6 | 28.7 ± 11.8 | 25.8 ± 9 | |
|  | 14–35 | 15–44 | 18–40 | - |
|  | 10 | 6 | 5 |  |
| Shelled Eggs | 19 ± 6.3 | 27.7 ± 8.4 | 24.2 ± 5.8 | |
|  | 12–29 | 14–41 | 15–31 | - |
|  | 5 | 7 | 5 |  |
| Egg Length (mm) | 12.3 ± 1.1 | 14.5 ± 0.9 | 13.8 ± 1.5 | |
|  | 9.8–14.6 | 11.8–16.2 | 10.9–16.8 | - |
|  | 96 | 188 | 115 |  |
| Egg Width (mm) | 8.2 ± 0.8 | 10.3 ± 0.7 | 9.9 ± 1.3 | |
|  | 6.7–9.9 | 8.2–12 | 7–12 | - |
|  | 96 | 188 | 89 |  |
| Egg Volume (cm<sup>3</sup>) | 440.3 ± 105.9 | 810.9 ± 120.6 | 755.4 ± 243.6 | |
|  | 256.2–682.5 | 470.5–1,076.8 | 297.6–1,226.7 | - |
|  | 96 | 188 | 89 |  |
| Clutch Volume (cm<sup>3</sup>) | 8,365.9 ± 4,156.5 | 22,240.6 ± 8,289.9 | 16,808.3 ± 7,205.5 | |
|  | 3,675.1–14,931 | 9,715.8–35,964.7 | 10,472.4–25,450.8 | - |
|  | 5 | 7 | 4 |  |
| Female SVL with Shelled Eggs | 73.9 ± 8.9 | 102 ± 4.4 | 95.9 ± 7.9 | |
|  | 62.2–86.1 | 97.1–107.5 | 84.3–104.6 | - |
|  | 5 | 7 | 5 |  |
| Female SVL with Yolked Ovarian Follicles | 68.3 ± 5.9 | 98.1 ± 2.9 | 96.4 ± 5.1 | |
|  | 61.6–77.6 | 94.3–101.6 | 89.8–103.7 | - |
|  | 10 | 6 | 5 |  |

### Shelled egg size

Size-corrected analyses detected no significant differences among populations in the means of egg length (*F*_2,15_ = 0.1, *P* = 0.903), egg width (*F*_2,15_ = 1.16, *P* = 0.346), or egg volume (*F*_2,15_ = 0.73, *P* = 0.503) (Table 1). Because egg length (R^2^ = 61.3%, *F*_1,15_ = 23.79, *P* < 0.001), egg width (R^2^ = 85.2%, *F*_1.15_ = 87.03, *P* < 0.001), and egg volume (R^2^ = 81.8%, *F*_1,15_ = 63.11, *P* < 0.001) were positively correlated with SVL, we removed the effects of body size by calculating residual scores from the separate regressions of each metric (and clutch size) on maternal SVL. Using the resulting residuals in separate regression analyses revealed a negative relationship between clutch size and egg sizes—there was an overall decrease in residual egg length, egg width, and egg volume with increasing residual clutch size (Fig. 5A–C).

**Fig. 5.**
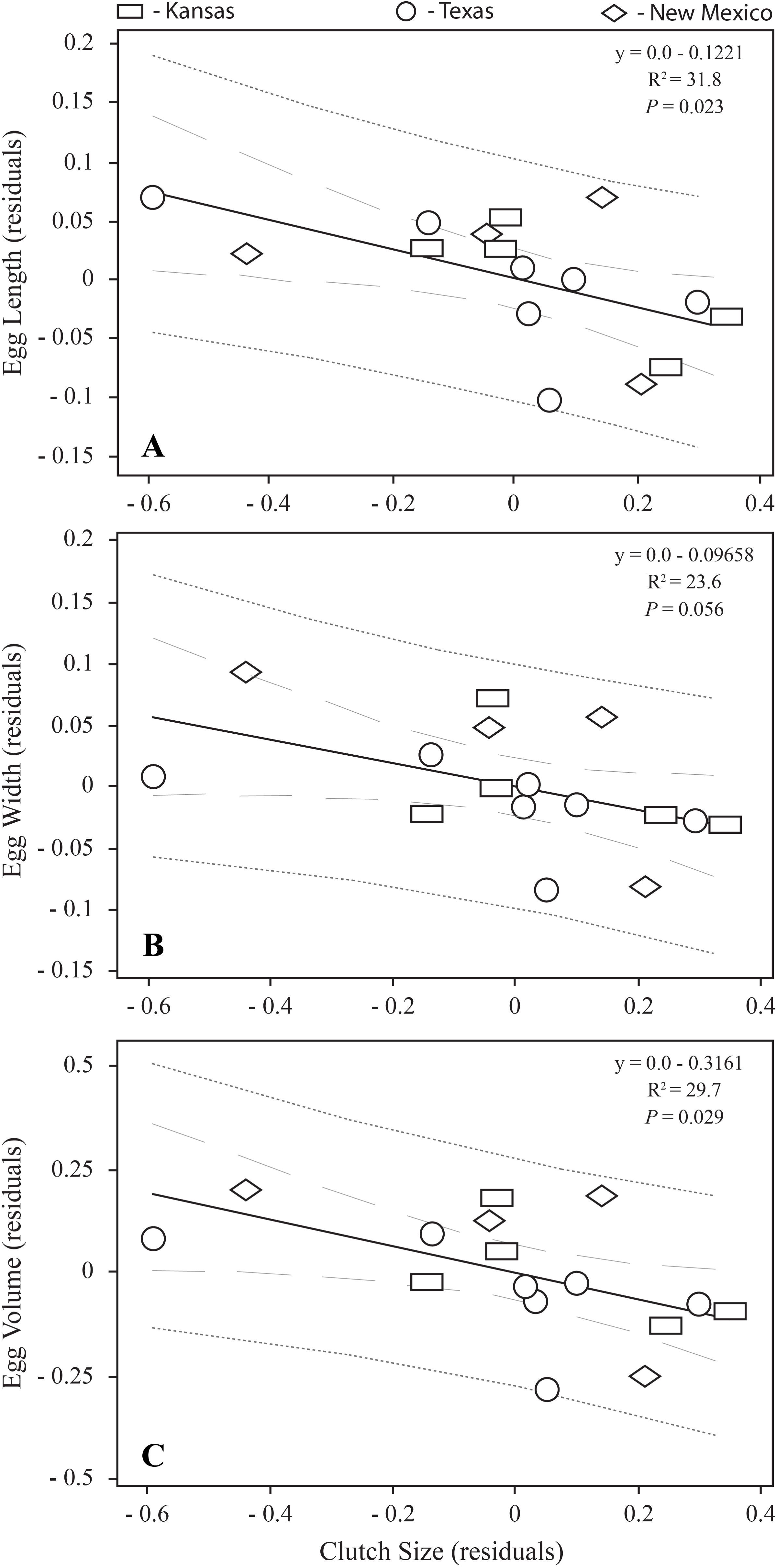
Tradeoff between egg size and clutch size after statistically removing the effects of maternal body size in three populations of the Texas horned lizard (*Phrynosoma cornutum*). (A) Residuals for egg length plotted against residual clutch size; (B) residuals for egg width plotted against residual clutch size; and (C) residuals for egg volume plotted against residual clutch size. The prediction interval is denoted by a dark-grey tightly spaced dashed line and the 95% confidence interval is denoted by a light-grey loosely spaced dashed line.

### Clutch frequency

We only detected luteal scars associated with expended oviductal eggs, and concurrent ovarian follicles measured 2–3 mm. We found a single female collected in June from Saline County, Kansas, that had 29 shelled eggs and 25 yolked ovarian follicles > 3 mm, suggesting that this individual had the potential to lay a second clutch that year given acceptable energetic or environmental conditions. These findings suggest that females from these three sites typically produce a single clutch annually, however, we could not rule out multiple clutch production in some individuals.

### Monthly activity

We detected activity based on pooled capture dates across locations. We found that the activity in the Texas horned lizard ranged from four to seven months. Months of capture were longest in Texas (March–October), followed by Kansas (April–October), New Mexico (April–September), and Mexico (June–October). For three of the locations, captures ended in October.

### Age at sexual maturity

The smallest juvenile we examined measured 22.6 mm SVL from Ellis County, Kansas, captured on 14 September 1963. Body-size cohorts were apparent in monthly distributions of body sizes for Kansas (Fig. 4G). We presume May and June eggs to have hatched within two months of oviposition. We do not know how much growth would have occurred from July and August hatching to our first juvenile cohort in September (22.6–29.2 mm). However, little growth appeared between fall and the following spring capture (26.4 mm). Fastest growth would have resulted in sexual maturity of males as early 12–13 months of age. Females and all other males would have reached sexual maturity during their second spring of life at an age of about 21–22 months of age (Fig. 4G). Too few data were available for the remaining three locations to estimate growth rates from monthly distributions of body size (Fig 4. H–I).

### Body size at sexual maturity

Body sizes of adult males followed a latitudinal trend of increasing size from north to south with respect to minimum, maximum, and mean values (Fig. 6; Table 1). Mean male SVL in Kansas was significantly smaller than in New Mexico, Texas, and Mexico (*F*_3,105_ = 161.19, *P* < 0.001). Likewise, body sizes of adult females followed a geographic trend of increasing size from north to south with respect to minimum and mean values. Mean female SVL in Kansas was significantly smaller than in New Mexico, Texas, and Mexico (*F*_3,79_ = 54.63, *P* < 0.001) (Fig. 6; Table 1).

**Fig. 6.**
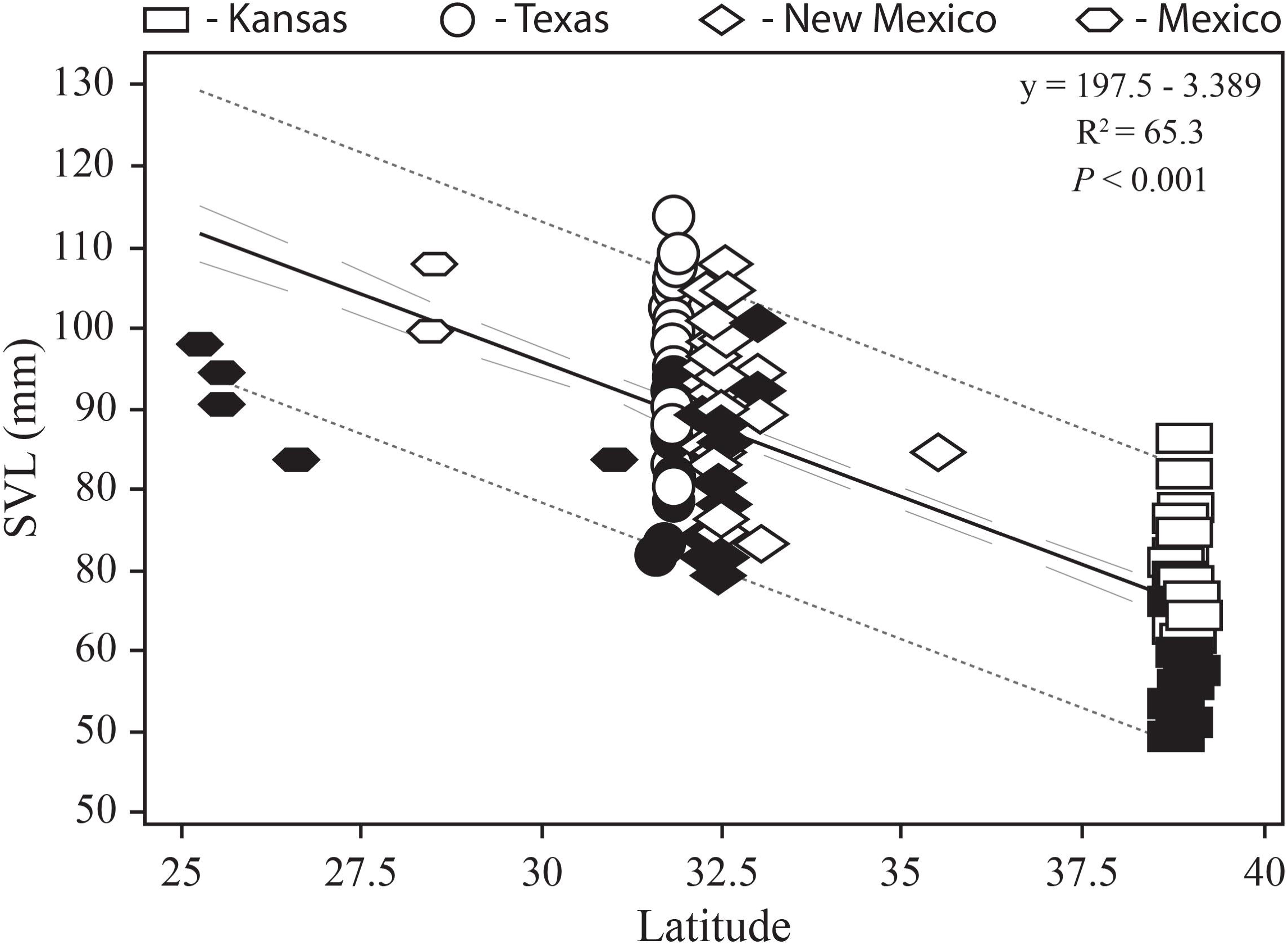
Body size regressed against latitude in four populations of the Texas horned lizard (*Phrynosoma cornutum*). The prediction interval is denoted by a dark-grey tightly spaced dashed line and the 95% confidence interval is denoted by a light-grey loosely spaced dashed line. Filled symbols represent males and open symbols females.

### Sexual dimorphism in adult body size

Sample sizes from three sites were large enough to test for differences in mean adult body sizes between the sexes. Mean body size of adult males was found to be significantly smaller than that of adult females from Kansas (*t* = −7.77, df = 63, *P* < 0.001), New Mexico (*t* = −3.06, df = 42, *P* = 0.004), and Texas (*t* = −5.76, df = 68, *P* < 0.001) (Table 1).

### Trends in multivariate life history

The pattern of differentiation among populations and latitudinal influences are presented in Fig. 7A–B. Relationships among populations were adequately represented in two dimensions judging by the magnitude of their respective eigenvalues; the first two dimensions explained 95% of the total variance (Fig. 7A). Three of the five variables loaded similarly on PC1, which overall described 65% of the variation in the data (Table 2). The first eigenvalue showed a moderate negative loading on clutch volume and clutch size, and moderate positive loadings on egg length, egg width, and egg volume. For PC2, which explained 30% of the variance, the second eigenvalue had the highest positive loading for clutch volume and relatively high for clutch size, and low positive loadings for egg width, egg volume, and egg length (Table 2).

**Fig. 7.**
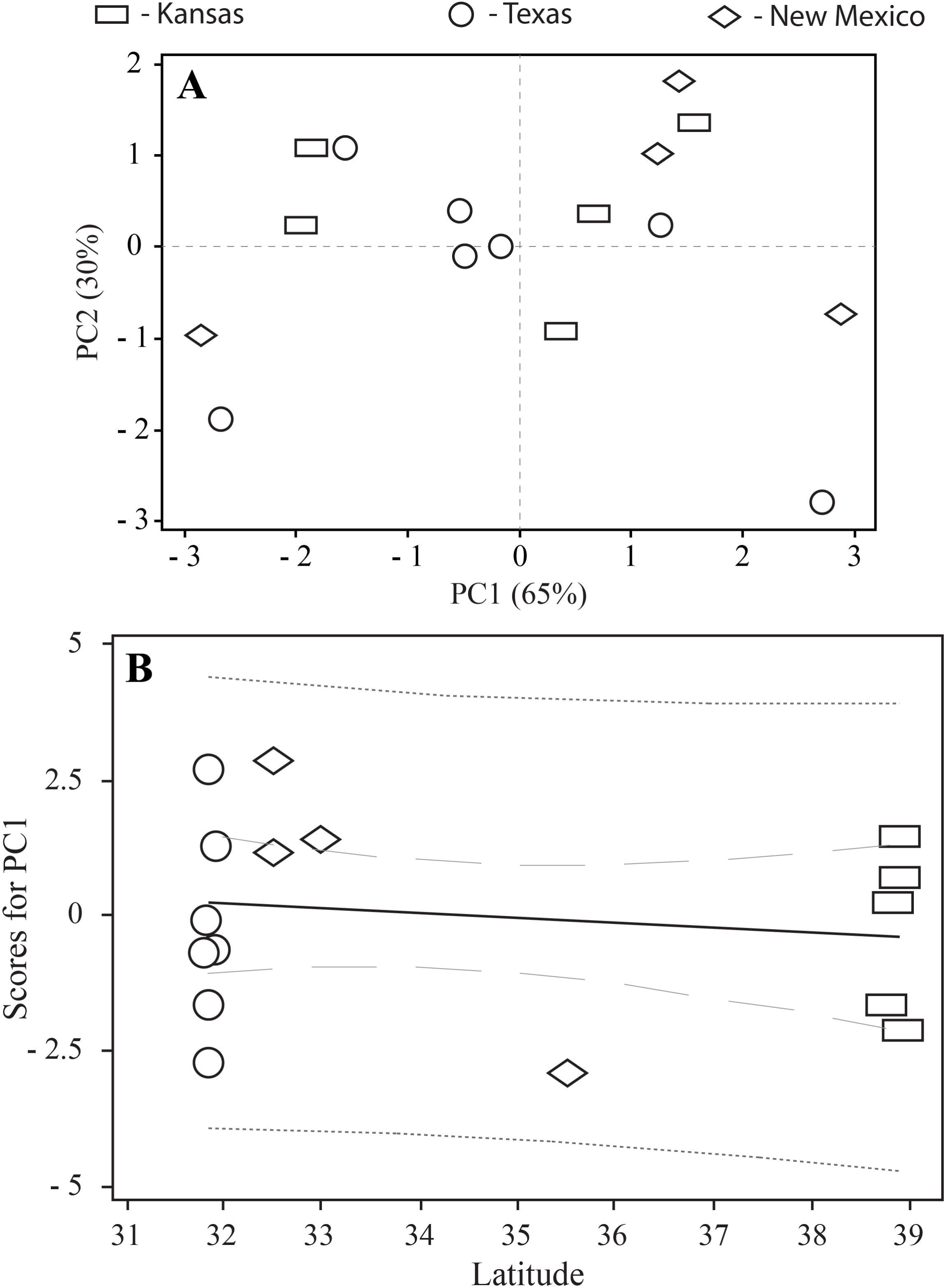
Multivariate life-history divergence for three populations of the Texas horned lizard (*Phrynosoma cornutum*). (A) Scatter plot of the first two principal components extracted from five life-history variables that were corrected for maternal body size. (B) Scores of principal components 1 (see Table 2 for loadings) plotted against latitude (y = 3.061 – 0.089x, R^2^ = 2.5%, *P* = 0.556).

**Table 2.** Principal components analysis comparing populations of the Texas horned lizard (*Phrynosoma cornutum*) with five life-history characters regressed against snout–vent length (SVL). Loadings, eigenvalues, variance, and cumulative variance are shown for the first two principal components. Values in bold represent the variables with the highest loadings.

|  | **Variable** | **PC1** | **PC2** |
| --- | --- | --- | --- |
| **1** | Clutch Size/SVL | – 0.442 | **0.486** |
| **2** | Egg Length/SVL | **0.496** | 0.172 |
| **3** | Egg Width/SVL | **0.486** | 0.318 |
| **4** | Egg Volume/SVL | **0.519** | 0.283 |
| **5** | Clutch Volume/SVL | – 0.231 | **0.743** |
|  | Eigenvalue | 3.259 | 1.483 |
|  | Proportion (variance) | 0.65 | 0.30 |
|  | Cumulative (variance) | 0.65 | 0.95 |

The majority of the variation in our data was described by PC1 and PC2 which were easily interpreted. Thus, we used the scores for these axes to test the hypothesis that latitudinal clines are derived from a co-adaptation of life-history traits. The scores on the first principal component showed a non-significant relationship with latitude (R^2^ = 2.5%, *F*_1,15_ = 0.36, *P* = 0.556) (Fig. 7B). The second principal component also exhibited a non-significant relationship with latitude (R^2^ = 4.7%, *F*_1,15_ = 0.69, *P* = 0.421). We did not find support for the hypothesis that the clutch measurements we included in this analysis covary with latitude.

## DISCUSSION

The life-history strategy of Texas horned lizards seems to have remained highly conserved as the species expanded its range northward after the last glaciation. The nesting season in Kansas is delayed by just a month compared to Texas, and in New Mexico the season spans from the beginning of that in Texas to the end of that in Kansas. The male gonadal cycle of maximal size in spring, minimal size in summer, and recrudescence in fall is generally consistent from 39°N to 25°N. Further, the overall pattern of monthly activity is nearly identical from Mexico to Kansas, and a two-year interval to reach maturity is also maintained across latitude. The similarity among populations is even more remarkable given the fact that drastic climatic differences exist among locations and our findings of adult body size differences. We did find pronounced differences in clutch sizes among Texas horned lizards, however. We detected latitudinal signatures in body sizes and consequently clutch sizes, but did not detect a strong influence of latitude in multivariate life history among populations. We will first discuss reasons as to why seasonal activity patterns and age at maturity are so conserved in Texas horned lizards and then we address latitudinal variation in life history and the implications of our results.

The nesting season of Texas horned lizards, for the most part, is restricted to a few summer months: May–June in Texas (Pianka and Parker, 1975) and Kansas (this study), May– July in southern Kansas (Gilver, 1922), central Texas (Ballinger, 1974), and southern New Mexico (this study), June–July in western Texas (this study), and June–August in the southwestern United States (Howard, 1974). Our results corroborate the evidence for a relatively short nesting season throughout its range in the United States, and, as did Gilver (1922) in southern Kansas, we found that the northernmost population exhibits a strikingly similar range. Our expectation that nesting activity patterns would vary latitudinally was based on temperature being of significant importance to ectotherm physiology (Huey, 1982). However, monthly activity and the nesting season seemed to exhibit only minor shifts in relation to temperature changes. Nesting and activity do not appear to simply be explained by temperature, at least in Kansas. Thus, some other factor that varies seasonally must underlie these behaviors. Lizards behaviorally thermoregulate to maintain activity levels necessary to meet seasonal needs such as reproduction (e.g., Adolph, 1990). Mating in the Texas horned lizard occurs largely in the spring and can extend into early summer: April–May in Kansas, April–June in Texas, and June–July in New Mexico. Increased activity during the mating season suggests that, following mating, the lizards become less active than thermal conditions would allow. This pattern presumably represents a tradeoff between activity and survival.

Characterized as delayed breeders, several species of horned lizards reach sexual maturity at two years of age (Pianka and Parker, 1975), including the Texas horned lizard (Ballinger, 1974; Howard, 1974; Endriss et al., 2007). Using our smallest individual (22.6 mm SVL), a two year-estimate of age at sexual maturity in northern Kansas does not differ from that of other studies, which is even more remarkable in light of marked differences in body sizes among populations. Juveniles and hatchling Texas horned lizards typically are found in the Fall across its range: August in central Texas (Ballinger, 1974), June–October from several sites along the US–Mexico border (Howard, 1974), late-July–August in central Oklahoma (Endriss et al., 2007), and late-August–September in Colorado (Montgomery and Mackessy, 2003). We do not know if all of the cohort of young individuals taken in September from northern Kansas represent hatchlings or a mix of both hatchlings and individuals hatched earlier in the season. The latter seems likely; based on likely hatching months, the earliest maturing individuals in northern Kansas would still not breed for the first time until nine months later. Our findings are inconsistent with Adolph and Porter’s (1996) model that populations from colder environments mature at a later age and at a larger size than populations in warmer areas (e.g., Wapstra et al., 2001). Alternatively, Texas horned lizards occupying colder, northern habitats reached sexual maturity at the same age as their counterparts from warmer, southern areas but at smaller sizes. Several other phrynosomatid lizards exhibit this same general trend (e.g., Parker and Pianka, 1975; Mathies and Andrews 1995), as does an Australian skink (Forsman and Shine, 1995).

From Colorado to Mexico, adult body sizes in Texas horned lizards of both sexes decreased from south to north in a latitudinal cline (Montgomery et al., 2003). We found a similar pattern of decreasing adult body size with increasing latitude. A latitudinal trend in body size is often attributed to differences in climate decreasing activity (Angilletta et al., 2004), environmental differences in food availability (Ashmole, 1963), or even day length (Rose and Lyon, 2013). Our results suggest that to maintain the highly conserved two-year constraint to reach sexual maturity, adult body sizes exhibit marked latitudinal differences dependent upon environmental variables. Indeed, we found that the most northern population and those in the middle of its range exhibit the same pattern of delayed maturity, but maturity was reached at a smaller size to compensate for reduced activity and likely reduced growth at high latitudes.

A significant departure in the life history of the Texas horned lizard was found in the clutch characteristics in our samples. Across its geographic range, the Texas horned lizard produces large clutches (Givler, 1922; Ballinger, 1974; Howard, 1974; Pianka and Parker, 1975; Vitt, 1977; Endriss et al., 2007), and our data corroborate that finding. A significant difference, however, exists among our sites, such that females from the northernmost site produced the smallest clutches. This finding also held true when female body size was held constant among our sites. Thus, selection seems to work against fecundity in Texas horned lizards at the northern edge of their range, placing them at a unique disadvantage of diminishing returns in energy investment to reproductive output. Ballinger (1974) found no evidence of geographic variation in clutch size, either latitudinally in Texas or longitudinally along the US–Mexico borderlands. The absence of a latitudinal trend in his study is understandable. We suspect that Ballinger’s (1974) finding is best explained by sampling of the species within the center of its geographic range, where clutch sizes are all similarly large.

Egg size in Texas horned lizards has not been examined previously with respect to clutch size. With female SVL held constant, we found that the relationships between clutch size and three measures of egg size were negative. The tradeoff between offspring number and egg size is mediated by a fecundity advantage of producing small offspring balanced against the survival advantage of large offspring. Although not measured for the Texas horned lizard, egg size is generally correlated with offspring size in lizards (Sinervo, 1990; Forsman and Shine, 1995). We found strong support for the prediction that clutch size is inversely related to egg size (i.e., offspring size) (Stewart, 1979). The amount of energy allocated to reproduction by females is divided between clutch size and egg size, and thus as clutch size increases egg size decreases. In fact, many oviparous squamates exhibit this tradeoff between the number and the size of offspring (e.g., Ford and Seigel, 1989).

Although a single clutch per year is produced in central Texas (Ballinger, 1974) and eastern Arizona (Vitt, 1977), multiple annual clutch production appears to more common than previously thought in Texas horned lizards. We could not entirely rule out the possibility of multiple clutches from our Texas and New Mexico samples and we found one female in Kansas that had shelled eggs and yolked ovarian follicles (> 3 mm) concurrently. Howard (1974) found evidence of double clutching in a sample of 21 females from four populations ranging longitudinally along the US–Mexico border. Wolf (2012) found that 6 of 9 radio-tracked females from central Oklahoma deposited two clutches in 2011. Further, Wolf et al. (2014) showed that nearly half of the females from central Oklahoma and about a third from south Texas produced two clutches annually. There remains the possibility that triple-clutching occurs in southern Texas populations (Burrow, 2000) based on observations of continual nesting throughout the active season (Wolf, 2012; Wolf et al., 2014). Texas horned lizards exhibit a slightly longer season of activity in south Texas (February–December with most activity April–August [Moeller et al., 2005]), which may contribute to additional egg-laying bouts. Sufficient time for the production of multiple clutches is feasible in south Texas. However, this interval does not explain multiple clutch production in Oklahoma. Mechanisms underlying multiple clutch production in northern populations of the Texas horned lizard are unclear, and may be related to a combination of favorable environmental triggers such as warmer temperatures and increased food intake. The effects of these variables on horned lizard fecundity are worth exploring.

From their respective study sites, Ballinger (1974) considered the Texas horned lizard to be K-selected for most traits (e.g., delayed maturity), yet r-selected for clutch size and juvenile survivorship, and Howard (1974) concluded that the species was also K-selected for late-maturity and r-selected to produce multiple large clutches annually. Reproductive characteristics, age at sexual maturity, and adult body sizes of the Texas horned lizard in our study included specimens from sites much farther north than examined by Howard (1974) and Ballinger (1974). Those specimens provided the material necessary to test the extent to which earlier findings concerning life history traits were applicable to this species and the genus. Our analyses show extreme conservatism in many life-history characteristics across its range, yet we detected a considerable extent of geographic variation in several direct measures of fecundity.

### Conclusions

The Texas horned lizard is a geographically widespread species (Price, 1990) whose categorization in life-history theory has been based on populations largely exclusive of the most northern and southern latitudes. Interestingly, even with samples from the northern edge of its range included, we found that the species generally conformed to Ballinger’s (1974) findings in Texas populations and Pianka and Parker’s (1975) genus-wide model. However, we found the smallest clutch sizes from populations in the northernmost locations and, importantly, a reduction in clutch size was not a simple consequence of body size variation among populations. Significant differences were still evident after we statistically controlled for the effects of maternal body size. Geographic variation in body size varies among many species of reptiles, and directions or absences of latitudinal signatures in this trait vary among them (e.g., Meshaka and Layne, 2015). In Texas horned lizards, the reproductive costs associated with living in northern Kansas is the most extreme (Shine, 1992). Unlike some of the Texas populations comprised of very large females that produce a single large clutch or multiple clutches each year, those in Kansas are restricted to a significantly smaller clutch with a potentially reduced likelihood of multiple clutch production. It remains to be tested as to what the latitudinal endpoint is for this ostensibly disadvantageous life-history response in the Texas horned lizard.

We are intrigued by the reduction in clutch size apart from female body size in northern Kansas because these populations persist at a distinct disadvantage with respect to fecundity. Further, we wonder what selective pressures drive the lower fecundity in the northern edge of its range, and we consider that this species at that latitude may be subjected to reduced opportunities for not only foraging on a bereft prey base and but also for photosynthesizing vitamin D_3_, a hormone necessary for calcium metabolism (Holick, 2010). Whitford and Bryant (1979) demonstrated irrefutably that Texas horned lizards are food limited and exhibit a pronounced preference for harvester ants (*Pogonomyrmex* spp.). Although the extent of this dietary dependence is subject to variation (e.g., Milne and Milne, 1950; Ramakrishnan et al., 2018), harvester ant densities and colony sizes are geographically variable (MacMahon et al., 2000; Warburg et al., 2017). In fact, harvester ant densities were found to be far greater at a central Texas site compared to a southern New Mexico site (Whiting et al., 1993). Also, the diversity of New World ants follows a latitudinal cline with fewer species at higher latitudes (Kaspari et al., 2003), and this pattern is consistent for seed-eating ants across the American Southwest (Davidson, 1977). At northern latitudes, lizard life histories are sensitive to both reductions in nutritional energy intake (Ballinger, 1977) and the availability of suitable thermal microclimates as they relate to the endogenous production of vitamin D_3_ by exposure to ultraviolet-B (290–315 nm)—a major component of nutritional quality for the Texas horned lizard (Ferguson et al., 2015) and crucial for lizard reproduction (Ferguson et al., 1996). The apparent disadvantage in fecundity we observed in northern populations of the Texas horned lizard may be related to compromised opportunities for vitamin D_3_ production from reduced aboveground activity. We suggest that not only a reduction in the extent of its preferred prey base—both diversity and abundance—but also decreased opportunities to produce vitamin D_3_ are linked to the diminished fecundity in Texas horned lizard populations at northern latitudes.

## ACKNOWLEDGMENTS

First and foremost, we dedicate this work to Royce E. Ballinger and the late C. Wayne Howard in recognition for their excellent contributions to this topic over 40 years ago. This study would not have been possible without the personal and professional kindnesses of Curtis J. Schmidt, Curator of vertebrate collections of the Sternberg Museum and we extend our most sincere gratitude to him.

## LITERATURE CITED

Adolph, S. C. 1990. Influence of behavioral thermoregulation on microhabitat use by two *Sceloporus* lizards. Ecology 71:315–327.

Adolph, S. C., and W. P. Porter. 1993. Temperature, activity, and lizard life histories. The American Naturalist 142:273–295.

Adolph, S. C., and W. P. Porter. 1996. Growth, seasonality, and lizard life histories: age and size at maturity. Oikos 77:267–278.

Ashton, K. G., and C. R. Feldman. 2003. Bergmann’s rule in nonavian reptiles: turtles follow it, lizards and snakes reverse it. Evolution 57:1151–1163.

Ashmole, N. P. 1963. The regulation of numbers of tropical oceanic birds. Ibis 103b:458–473.

Atkinson, D. 1994. Temperature and organism size: a biological law for ectotherms? Advances in Ecological Research 25:1–58.

Ballinger, R. E. 1974. Reproduction of the Texas horned lizard, *Phrynosoma cornutum*. Herpetologica 30:321–327.

Ballinger, R. E. 1977. Reproductive strategies: food availability as a source of proximal variation in a lizard. Ecology 58:628–635.

Blouin-Demers, G., K. A. Prior, and P. J. Weatherhead. 2002. Comparative demography of black rat snakes (*Elaphe obsoleta*) in Ontario and Maryland. Journal of Zoology 256:1–10.

Burrow, A. L. 2000. The effect of prescribed burning and grazing on the threatened Texas horned lizard (*Phrynosoma cornutum*) in the western Rio Grande plains. Unpubl. M.S. thesis, Oklahoma State University, Stillwater, Oklahoma, USA.

Davidson, D. W. 1977. Species diversity and community organization in desert seed□eating ants. Ecology 58:711–724.

Endriss, D. A., E. C. Hellgren, S. F. Fox, and R. W. Moody. 2007. Demography of an urban population of the Texas horned lizard (*Phrynosoma cornutum*) in in central Oklahoma. Herpetologica 63:320–331.

Ferguson, G. W., J. R., Jones, W. H. Gehrmann, S. H. Hammack, L. G. Talent, R. D. Hudson, E. S. Dierenfeld, M. P. Fitzpatrick, F. L. Frye, M. F. Holick, and T.C. Chen. 1996. Indoor husbandry of the panther chameleon *Chamaeleo* [*Furcifer*] *pardalis*: effects of dietary vitamins A and D and ultraviolet irradiation on pathology and life-history traits. Zoo Biology 15:279–299.

Ferguson, G. W., W. H. Gehrmann, A. M. Brinker, G. C. Kroh, and D. C. Ruthven III. 2015. Natural ultraviolet-b exposure of the Texas Horned Lizard (*Phrynosoma cornutum*) at a North Texas Wildlife Refuge. The Southwestern Naturalist 60:231–239.

Fitch, H. S. 1985. Variation in clutch and litter size in New World reptiles. University of Kansas Museum of Natural History Miscellaneous Publications 76:1–76.

Ford, N. B., and R. A. Seigel. 1989. Relationships among body size, clutch size, and egg size in three species of oviparous snakes. Herpetologica 45:75–83.

Forsman, A., and R. Shine. 1995. Parallel geographic variation in body shape and reproductive life history within the Australian scincid lizard *Lampropholis delicata*. Functional Ecology 9:818–828.

Givler, J. P. 1922. Notes on the oecology and life-history of the Texas horned lizard, *Phrynosoma cornutum*. Journal of the Elisha Mitchell Scientific Society 37:130–137.

Hammerson, G. A. 2007. Phrynosoma cornutum. The IUCN Red List of Threatened Species 2007: e.T64072A12741535. http://www.iucnredlist.org/details/6407270 (Accessed 4 October 2017).

Holick, M. F. (ed.). 2010. Vitamin D: Physiology, Molecular Biology, and Clinical Application. Second edition. Humana Press, New York, USA.

Howard, C. W. 1974. Comparative reproductive ecology of horned lizards (Genus *Phrynosoma*) in southwestern United States and northern Mexico. Journal of the Arizona Academy of Science 9:108–116.

Huey, R. B. 1982. Temperature, physiology, and the ecology of reptiles, p. 25–91. *In*: Biology of the Reptilia, Physiology (C), Volume 12. C. Gans, and F. H. Pough (eds.). Academic Press, London, UK.

Huey, R. B., and D. Berrigan. 2001. Temperature, demography, and ectotherm fitness. The American Naturalist 158:204–210.

Kaspari, M., M. Yuan, and L. Alonso. 2003. Spatial grain and the causes of regional diversity gradients in ants. The American Naturalist 161:459–477.

King, R. B. 2000. Analyzing the relationship between clutch size and female body size in reptiles. Journal of Herpetology 34:148–150.

Laurila, A., B. Lindgren, and A. T. Laugen. 2008. Antipredator defenses along a latitudinal gradient in *Rana temporaria*. Ecology 89:1399–1413.

Laugen, A. T., A. Laurila, K. Räsänen, and J. Merilä. 2003. Latitudinal countergradient variation in the common frog (*Rana temporaria*) development rates–evidence for local adaptation. Journal of Evolutionary Biology 16:996–1005.

MacMahon, J. A., J F. Mull, and T. O. Crist. 2000. Harvester ants (*Pogonomyrmex* spp.): their community and ecosystem influences. Annual Review of Ecology and Systematics 31:265–291.

Mathies, T., and R. M. Andrews. 1995. Thermal and reproductive biology of high and low elevation populations of the lizard *Sceloporus scalaris*: implications for the evolution of viviparity. Oecologia 104:101–111.

Mayhew, W. W. 1963. Reproduction in the granite spiny lizard, *Sceloporus orcutti*. Copeia 1963:144–152.

McGarigal, K., S. Cushman, and S. Stafford. 2000. Multivariate Statistics for Wildlife and Ecology Research. Springer, New York, USA.

Meshaka, W. E. Jr., and J. N. Layne. 2015. The Herpetology of southern Florida. Herpetological Conservation and Biology 10(Monograph 5):1–353.

Milne, L. J., and M. J. Milne. 1950. Notes on the behavior of horned toads. The American Midland Naturalist 44:720–741.

Moeller, B. A., E. C. Hellgren, D. C. Ruthven III, R. T. Kazmaier, and D. R. Synatzske. 2005. Temporal differences in activity patterns of male and female Texas horned lizards (*Phrynosoma cornutum*) in southern Texas. Journal of Herpetology 39:336–339.

Montgomery, C. E., and S. P. Mackessy. 2003. Natural history of the Texas horned lizard, *Phrynosoma cornutum* (Phrynosomatidae), in southeastern Colorado. The Southwestern Naturalist 48:111–118.

Montgomery, C. E., S. P. Mackessy, and J. C. Moore. 2003. Body size variation in the Texas horned lizard, *Phrynosoma cornutum*, from central Mexico to Colorado. Journal of Herpetology 37:550–553.

Olalla-Tárraga, M. Á., M. Á. Rodríguez, and B. A. Hawkins. 2006. Broad□scale patterns of body size in squamate reptiles of Europe and North America. Journal of Biogeography 33:781–793.

Packard, G. C., and T. J. Boardman. 1999. The use of percentages and size-specific indices to normalize physiological data for variation in body size: wasted time, wasted effort? Comparative Biochemistry and Physiology Part A: Molecular Integrative Physiology 122:37–44.

Parker, W. S., and E. R. Pianka. 1975. Comparative ecology of populations of the lizard *Uta stansburiana*. Copeia 1975:615–632.

Pianka, E. R. 1970. On *r*- and *K*-selection. The American Naturalist 104:592–597.

Pianka, E. R., and W. S. Parker. 1975. Ecology of horned lizards: a review with special reference to *Phrynosoma platyrhinos*. Copeia 1975:141–162.

Price, A. H. 1990. Phrynosoma cornutum. Catalogue of American Amphibians and Reptiles 469: 1–7.

Ramakrishnan, S., A. J. Wolf, E. C. Hellgren, R. W. Moody, and V. Bogosian III. 2018. Diet selection by a lizard ant-specialist in an urban system bereft of preferred prey. Journal of Herpetology 52:79–85.

Rose, A. P., and B. E. Lyon. 2013. Day length, reproductive effort, and the avian latitudinal clutch size gradient. Ecology 94:1327–1337.

Shine, R. 1992. “Costs” of reproduction in reptiles. Oecologia 46:92–100.

Sinervo, B. 1990. The evolution of maternal investment in lizards: an experimental and comparative analysis of egg size and its effects on offspring performance. Evolution 44:279–294.

Sperry, J. H., G. Blouin-Demers, G. L. Carfagno, and P. J. Weatherhead. 2010. Latitudinal variation in seasonal activity and mortality in ratsnakes (*Elaphe obsoleta*). Ecology 91:1860–1866.

Stearns, S. C. 1977. The evolution of life history traits: a critique of the theory and a review of the data. Annual Review of Ecology and Systematics 8:145–171.

Stewart, J. R. 1979. The balance between number and size of young in the live-bearing lizard *Gerrhonotus coeruleus*. Herpetologica 35:342–350.

Tinkle, D. W., H. M. Wilbur, and S. G. Tilley. 1970. Evolutionary strategies in lizard reproduction. Evolution 24:55–74.

Tsuji, J. S. 1988. Thermal acclimation of metabolism in *Sceloporus* lizards from different latitudes. Physiological Zoology 61:241–253.

Van Noordwijk, A. J., and G. de Jong. 1986. Acquisition and allocation of resources: their influence on variation in life history tactics. The American Naturalist 128:137–142.

Vitt, L. J. 1977. Observations on clutch and egg size and evidence for multiple clutches in some lizards of southwestern United States. Herpetologica 33:333–338.

Warburg, I., W. G. Whitford, and Y. Steinberger. 2017. Colony size and foraging strategies in desert seed harvester ants. Journal of Arid Environments 145:18–23.

Wapstra, E., R. Swain, and J. M. O’Reilly. 2001. Geographic variation in age and size at maturity in a small Australian viviparous skink. Copeia 2001:646–655.

Whitford, W. G., and M. Bryant. 1979. Behavior of a predator and its prey: the horned lizard (*Phrynosoma cornutum*) and harvester ants (*Pogonomyrmex* spp.). Ecology 60:686–694.

Whiting, M. J., J. R. Dixon, and R. C. Murray. 1993. Spatial distribution of a population of Texas horned lizards (*Phrynosoma cornutum*: Phrynosomatidae) relative to habitat and prey. The Southwestern Naturalist 38:150–154.

Wolf, A. J. 2012. Spatial and demographic ecology of Texas horned lizards within a conservation framework. Unpubl. M.S. thesis, Southern Illinois University at Carbondale, Carbondale, IL, USA.

Wolf, A. J., E. C. Hellgren, E. M. Schauber, V. Bogosian, R. T. Kazmaier, D. C. Ruthven, and R. W. Moody. 2014. Variation in vital-rate sensitivity between populations of Texas horned lizards. Population Ecology 56:619–631.

